# Rational Development of Recombinant ELP Bolaamphiphiles for the Controlled Construction of Multifunctionalized Globular Protein Vesicles

**DOI:** 10.1101/2025.06.23.660353

**Authors:** Luis E. Lopez de la Maza, Bornita Deb, Jackson Powers, Ariory Hood, Yeongseon Jang, Carl A. Denard

## Abstract

Synthetic biology has enabled the development of new strategies for creating artificial cells that can sense and respond to external stimuli. This study introduces the bottom-up construction of globular protein vesicles (GPVs) that incorporate elastin-like peptide (ELP) bolaamphiphiles as transmembrane components. To enable this strategy, we devised a Golden Gate-based cloning strategy to streamline the design, expression, and purification of ELP bolaamphiphiles. Three ELP bolamphiphiles with varying structural complexity were developed, incorporating fluorescent proteins to facilitate visualization and characterization. The self-assembly of these bolaamphiphiles into GPVs was optimized by varying the molar ratios of recombinant building blocks. Structural characterization confirmed vesicle formation, dynamic light scattering analysis revealed size distributions dependent on modular complexity, and atomic force microscopy demonstrated that the vesicles exhibited MPa-range Young’s moduli, indicative of high mechanical robustness. Our findings demonstrate that multifunctional ELP bolaamphiphiles can be incorporated into GPVs, enabling modular vesicle engineering. This work provides a foundation for designing synthetic cells with customizable bi-functionalities and modularity, advancing compartmentalized systems.

## Introduction

Globular protein vesicles (GPVs), comprised entirely of recombinant protein fusions, have emerged as a highly versatile platform for constructing synthetic cellular systems.^1^ Park and Champion first demonstrated the thermally triggered formation of GPVs using fusion proteins mCherry-ZE (mCherry fused to glutamic acid-rich leucine zipper) and ZR-ELP (arginine-rich leucine zipper fused to elastin-like peptide), which form amphiphilic complexes through strong leucine zipper heterodimerization (*K*_*d*_∼10^-15^M) and lower critical solution transition (LCST) behavior of ELP.^2^ GPVs offer unique advantages over traditional polymersomes, liposomes, and proteoliposomes, including high biocompatibility, ease of functionalization, and mild, non-denaturing assembly conditions, making them attractive candidates for diverse applications in synthetic biology, such as vaccines and drug delivery.^3–12^ This platform has since been expanded to incorporate various globular proteins such as human carbonic anhydrase II (HCA), human glucokinase (HGK), and *E. coli* malate synthase G (MSG), further emphasizing its flexibility and potential.^4^ Our group has recently demonstrated that recombinant protein building blocks can be engineered to endow GPVs with sensing capabilities by integrating sensory domains such as FK506-binding protein (FKBP) and FKBP-rapamycin binding domain (FRB) into the vesicle membrane.^6^ This capability, combined with the vesicles’ tunable permeability, fluidic membrane nature, and mechanical robustness, makes GPVs a promising platform for mimicking complex cellular behaviors in a simplified, programmable format.^5,7,13^

A key motivation for advancing GPV technology is its potential to mimic transmembrane signal transduction (TST)—a hallmark of natural cellular systems that allows cells to sense and respond to external stimuli.^11–15^ Most natural and all synthetic TST proteins are composed of three parts: extracellular, transmembrane, and cytoplasmic domains.^19–27^ From a design point of view, such TST proteins are bolaamphiphilic, where two mostly hydrophilic protein arms flank a hydrophobic transmembrane domain. As it relates to GPVs, a similar architecture could be modeled as an ELP bolaamphiphile, i.e., a hydrophobic ELP flanked by hydrophilic globular domains. Above their transition temperature, ELP-based bolaamphiphiles can adopt conformations suitable for forming planar or spherically curved monolayer membranes, thereby allowing spatially separated sensing and actuation domains to straddle the membrane.^28^ This architecture could pave the way for modular TST systems embedded entirely within protein-based vesicle membranes.

In this work, we demonstrate for the first time the integration of recombinant ELP bolaamphiphiles into GPV membranes by building GPVs with three distinct ELP bolaamphiphilic architectures. ELP bolaamphiphiles pose intrinsic challenges during cloning, assembly and purification, due to the repetitive nature of ELP-coding sequences and restrictive protein purification strategies.^29^ Additionally, achieving site-specific post-translational functionalization of such constructs requires precise control mechanisms. To enhance modularity and ease of construction, we leveraged SpyCatcher/SpyTag technology, which enables fast, irreversible, and site-specific conjugation of protein domains to ELPs.^30^ The spontaneous intermolecular amide bond formation in SpyCatcher/SpyTag binding is compatible with a bottom-up construction of GPVs as it avoids the need for harsh chemicals or external ligation enzymes (e.g., sortases) and can be genetically encoded with minimal effort.^31–33^ SpyCatcher/SpyTag bioconjugation has also been applied in other ELP-based systems for modular functionalization of protein assemblies.^34–37^

To complement site-specific bioconjugation, we designed a Golden Gate-based cloning strategy^38^ that enables efficient colony screening, expression, and purification of ELP bolaamphiphiles. Using these strategies, three ELP bolamphiphiles with varying structural complexity were developed, incorporating fluorescent proteins to facilitate visualization and characterization. The self-assembly of these bolaamphiphiles into GPVs was optimized by varying the molar ratios of recombinant building blocks. Fluorescence imaging and structural characterization demonstrated that multifunctional ELP bolaamphiphiles can be stably incorporated into GPVs. Taken together, this study lays the foundation for building GPVs with cell-like sensor-actuator functions from multifunctionalized ELP bolaamphiphiles. This work also enables us to systematically investigate how the globular domain composition affects the self-assembly behavior and mechanical properties of GPV membranes.

## Results and Discussion

### Rational design for multifunctionalized GPVs leverages site-specific bioconjugation and seamless cloning

The GPV architecture that has been extensively reported is composed of two fusion proteins, a ZR-ELP and a water-soluble globular protein-ZE (i.e., mCherry-ZE), which combine to form an amphiphilic complex via a ZE/ZR interaction at temperatures above ELP LCST.^2^ As a result, protein solutions with a molar ratio (χ) of ZE to ZR in the range of 0.005 and 0.1 undergo a phase transition from soluble to self-assembled GPVs.^2,4–6,39,40^ We recently reported that hydrophilic fusion proteins containing an N-terminal receptor domain (i.e., FRB-mCherry-ZE and FKBP-GFP-ZE) can form sensory GPVs with ZR-ELP through thermally induced phase transition.

We aimed to extend this design to generate asymmetric bolaamphiphilic fusion proteins, in the form mCherry-ZE/ZR-ELP-globular protein, where an additional hydrophilic protein is appended to the C-termini of ELP. However, producing a recombinant ZR-ELP-globular protein is impractical because ZR fusion proteins produced in *E. coli* are required to be purified under denaturing conditions, limiting the globular proteins that could be tethered to the C-terminus of the ZR-ELP.^41,42^

To address this challenge, we leveraged the modularity of the GPV platform and explored strategies to decouple ZR purification from C-terminal ELP fusions. On the one hand, we incorporated the SpyCatcher/SpyTag technologies to build two ELP bolaamphiphiles: mCherry-ZE/ZR-SpyTag(SpT)-SpyCatcher (SpC)-ELP-GFP (BB1) and mCherry-ZE-SpT-SpC-ELP-GFP (BB2). In addition, a bolaamphiphilic mCherry-ELP-GFP devoid of a ZE/ZR interaction was also constructed. Thus, we generated a series of recombinant protein parts (Figure 1A) that could be pre-assembled and subsequently mixed with ZR-ELP to form GPVs. For instance, a ZR-SpyTag, which is purified under denaturing conditions, can be coupled with a SpyCatcher-ELP-GFP, which was purified under native conditions, to form ZR-SpyTag/SpyCatcher-ELP-GFP. The subsequent addition of mCherry-ZE generates an mCherry-ZE/ZR-SpyTag/SpyCatcher-ELP-GFP. Conversely, adding the SpyTag to the C-terminus of the mCherry-

**Figure 1.**
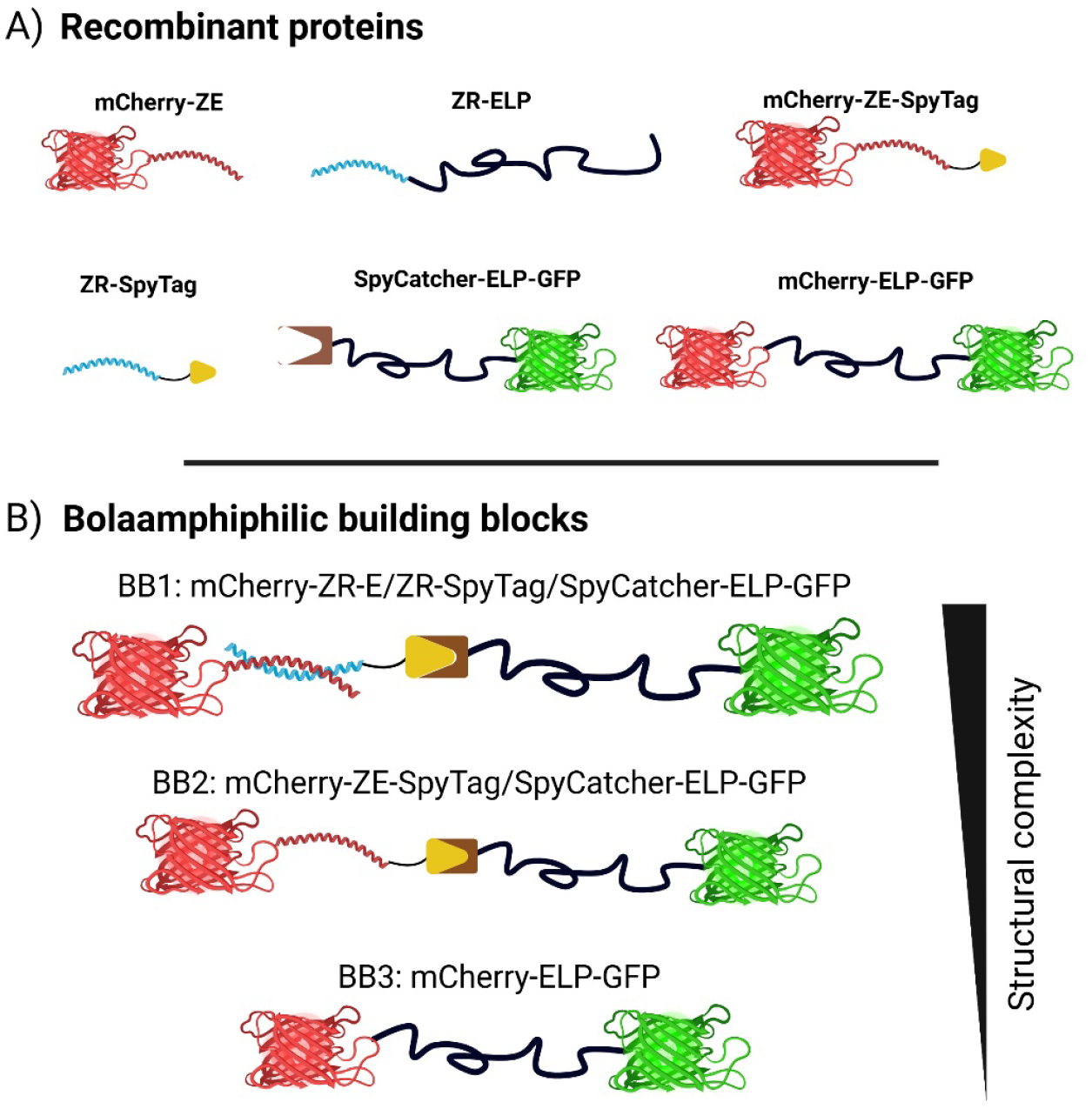
Molecular design of bolaamphiphilic building blocks for GPV assembly. (A) Schematic of modular recombinant fusion proteins used in this study. (B) Three bolaamphiphilic building blocks of varying modular complexity: BB1 (mCherry-ZE/ZR- SpyTag/SpyCatcher-ELP-GFP), BB2 (mCherry-ZE-SpyTag/SpyCatcher-ELP-GFP), and BB3 (mCherry-ELP-GFP). These constructs enable hierarchical assembly and membrane incorporation into GPVs.

ZE head domain allowed its conjugation to SpyCatcher-ELP-GFP, resulting in a bolaamphiphile devoid of a ZE/ZR interaction. Lastly, we completely bypassed both non-covalent (ZE/ZR) and covalent (SpyCatcher/SpyTag) interactions and generated an mCherry-ELP-GFP bolaamphiphile. These three types of bolaamphiphilic designs with varying levels of complexity enabled us to explore novel GPV architectures, thereby providing fundamental insights into the self-assembly requirements (Figure 1).

### Cloning, purification, and characterization of ELP bolaamphiphile parts

To build ELP bolaamphiphiles, we developed a Golden Gate cloning approach^38^ that incorporates several benefits: (i) cloning of ELPs with various lengths and compositions; (ii) N- and C-terminal addition to ELP; (iii) and an expression cloning approach, allowing the user to clone, verify expression, and perform protein purification in the same strain (Figure 2). pQE9 was selected as the template vector for the receiving Golden Gate plasmid. Two changes were made: (i) a silent mutation to delete the BsaI cut-site in the Ampicillin gene, and the insertion of a 10x His tag, a SpyCatcher or an mCherry gene, and two BsaI cut-sites downstream of the T5 promoter. Limited by the maximum ELP fragment length synthesizable by Twist Biosciences (9 repeats), three fragments are needed to assemble a 25-repeat ELP gene. Therefore, taking advantage of codon degeneracies with the VPGXG repeat of ELPs (where X is occupied with 80% Val and 20% Phe), we designed four unique overhangs that enable in-frame GG assembly of ELP fragments followed by a 3’ globular protein of choice (in our case, GFP). We also prepared two BsaI-containing receiver plasmids encoding N-terminal globular proteins, SpyCatcher and mCherry. Overall, this approach allowed us to rapidly clone and purify two ELP bolaamphiphiles, namely a SpyCatcher-ELP-GFP and an mCherry-ELP-GFP (Figure 2B, S1, S2, and Table S1). Meanwhile, we designed a plasmid to purify ZR-SpyTag, which occurs under denaturing conditions, as previously described for ZR-ELP (Table S2).^2,40^ Lastly, we generated an mCherry-ZE-SpyTag. Together with the traditional mCherry-ZE and ZR-ELP fusions, these proteins allow us to construct bolaamphiphilic ELPs, named building blocks (BB) 1, 2, and 3, and to test their assembly into GPVs of varying complexities.

**Figure 2.**
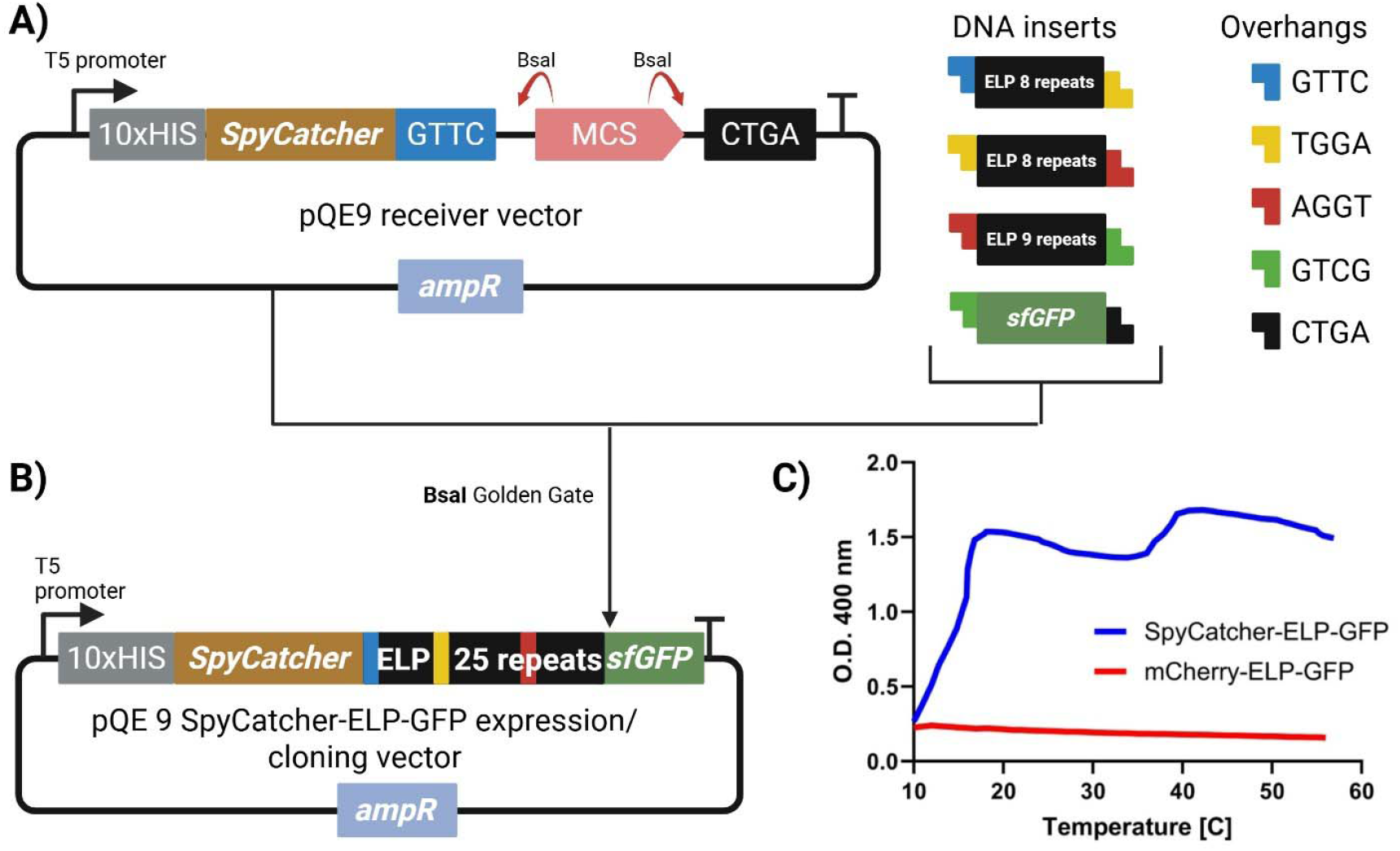
Golden Gate cloning workflow for constructing ELP bolaamphiphiles. (A) Modified pQE9 plasmid containing BsaI cut sites used as the assembly backbone for Golden Gate assembly and DNA fragments encoding ELP and GFP insert sequences. This approach allows efficient modular cloning of ELP-based bolaamphiphiles. (B) Assembly of SpyCatcher-ELP-GFP via Golden Gate. (C) Thermoresponsive phase transition of SpyCatcher-ELP-GFP and mCherry-ELP-GFP fusion proteins. Turbidity profiles were obtained by monitoring absorbance at 4001⍰m for 50⍰µM protein solutions in the presence of 1⍰M NaCl during a linear temperature ramp from 10⍰°C to 60⍰°C at 1⍰°C/min. The lower critical solution temperature (LCST) transition was defined as the inflection point of the absorbance curve.

We investigated whether a hydrophobic ELP bolaamphiphile would still undergo phase transition to form self-assembled GPVs by thermal triggers. We characterized the phase transition of SpyCatcher-ELP-GFP by monitoring changes in the optical density of protein solutions at 400 nm as a function of temperature. The transition temperature (T□), defined as the temperature at which the slope of the turbidity curve is maximized,^40,43^ was determined to be approximately 17 °C for a 50 μM solution of SpyCatcher-ELP-GFP in 1 M NaCl (Figure 2C). In a similar setup, ZR-ELP solutions (60 μM) have previously been shown to exhibit LCST behavior over the range of approximately 17-25 °C.39 Our observation is consistent with the expected LCST behavior of ELPs, wherein increased ionic strength promotes hydrophobic interactions and facilitates phase separation at lower temperatures.44,45 In contrast, a solution of mCherry-ELP-GFP under identical conditions (50 μM, 1 M NaCl) exhibited no significant turbidity changes across the temperature range tested (7–60 °C), suggesting a suppression or complete inhibition of the ELP phase transition (Figure 2C). This lack of observable transition may be attributed to enhanced hydrophilicity and steric effects introduced by the hydrophilic mCherry and GFP domains, which could interfere with ELP chain collapse or aggregation between them.^46^ These findings underscore the importance of fusion protein architecture in modulating ELP thermoresponsive behavior.^47^

### Highly modular, amphiphilic building blocks are incorporated in the vesicle membrane: BB1

Having built the various protein parts, we aimed to test their ability to form GPVs, starting with BB1 (mCherry-ZE/ZR-SpyTag/SpyCatcher-ELP-GFP). BB1 relies on two interactions: (i) the isopeptide bond formation between ZR-SpyTag and SpyCatcher-ELP-GFP and (ii) the heterodimerization of mCherry-ZE and ZR-SpyTag. This is the most complex design, as it preserves the ZE/ZR interaction previously reported in GPV studies.^10^ A stepwise assembly protocol was followed for GPV assembly containing BB1. First, ZR-SpyTag and SpyCatcher-ELP-GFP were ligated equimolarly (25 uM) at room temperature for 1 hour (Figure S3). The resulting ZR-SpyTag-SpyCatcher-ELP-GFP bolaamphiphile was incubated on ice with mCherry-ZE for 15 minutes in equimolar quantities to form the multifunctional building block mCherry-ZE/ZR-SpyTag-SpyCatcher-ELP-GFP. Finally, BB1 and ZR-ELP were mixed on ice for 15 minutes in the presence of 1 M NaCl to a final volume of 50 μL, at χ=0.01 and 0.05.

We observed the formation of GPVs under both conditions (Figure 3). At χ=0.01, isolated GPVs predominated in the sample, with large clusters of fully assembled GPVs also present (Figure 3A and S4A). This is not surprising because vesicle aggregation has been reported in the context of eGFP-ZE vesicles due to the high tendency of eGFP to form dimers.^39,48,49^ In the case of our bolaamphiphile designs, two possible causes for aggregation include: (i) excluded volume effects introduced by the SpyCatcher-SpyTag conjugate in the transmembrane domain and (ii) GFP dimerization at the vesicle membrane.^39,50^

**Figure 3.**
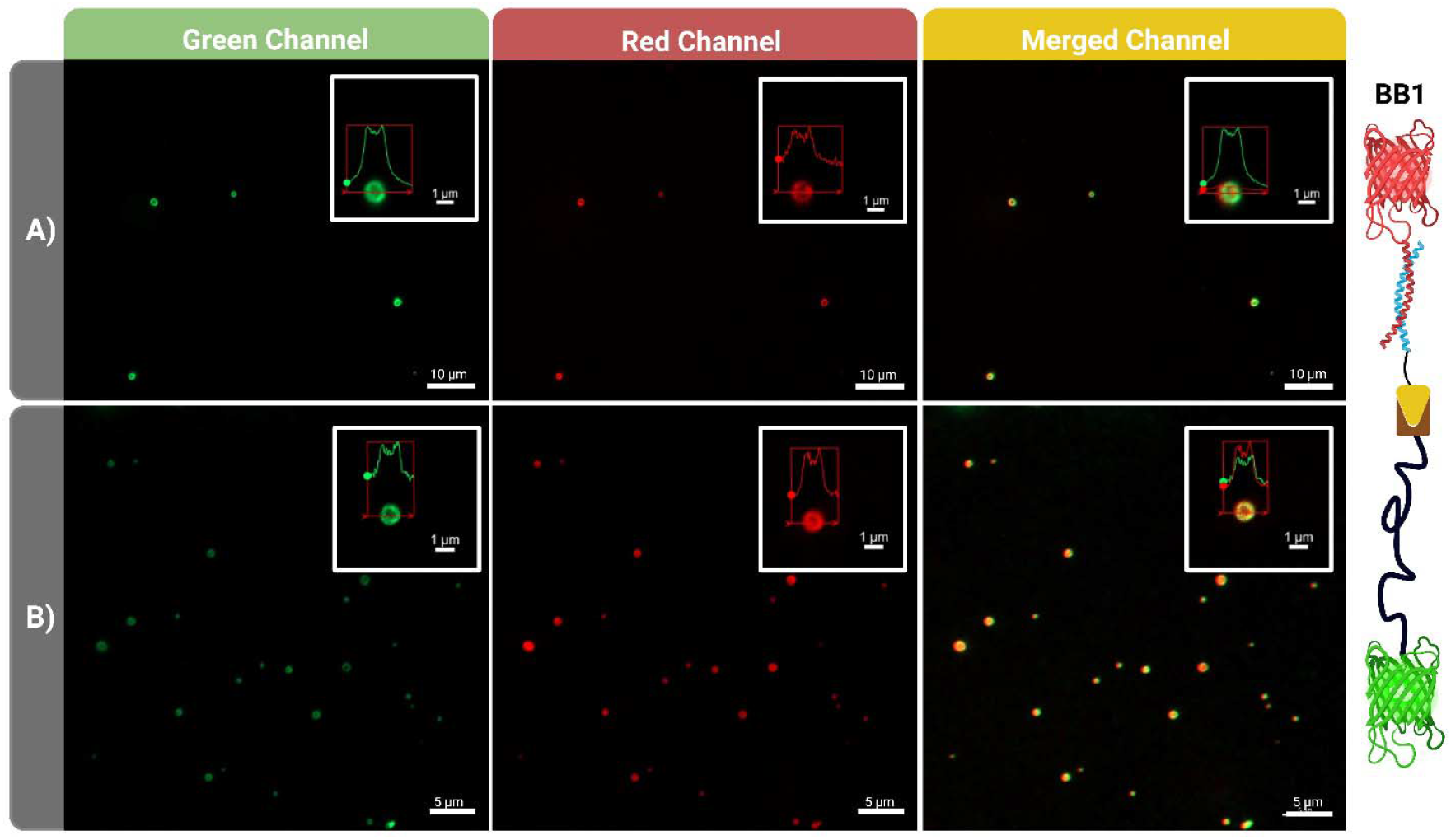
Epifluorescence micrographs of BB1-derived GPVs assembled with ZR-ELP. Vesicles were formed by incubating BB1 with ZR-ELP (60 μM) in 1 M NaCl for 1 hour at 25⍰°C. (A) χ = 0.01 molar ratio (BB1:ZR-ELP), (B) χ = 0.05.

At χ = 0.05, we observed a higher density of isolated GPVs relative to χ = 0.01, likely due to increased availability of BB1 (Figure 3B and S4B). However, vesicle clusters and coacervates were also present. The presence of coacervates suggests that for a given ion strength, temperature, and ELP concentration, there is a BB1 disorder-order transition concentration threshold. Shin *et al*. observed that the vesicle-to-coacervate transition occurs at specific molar ratios, even under constant ionic strength and temperature, suggesting that a compositional threshold governs protein phase behavior.^39^ This supports the notion that BB1 has a disorder–order transition concentration, which may also explain localized aggregation on the GPV membrane (Figure 3 and S4). Fluorescence imaging confirmed the successful co-localization of mCherry and GFP on the membrane of the assembled vesicles, highlighting the potential for membrane functionalization with multiple domains. The apparent structural complexity and organization of the membrane warrant future investigation using higher-resolution techniques.

### Reducing structural complexity of bolaamphiphilic GPV building blocks: BB2

BB2 (mCherry-ZE-SpyTag-SpyCatcher-ELP-GFP) simplifies the design of the building blocks to two units that come together spontaneously and irreversibly at room temperature. Globular proteins with a C-terminal SpyTag fusion and the SpyCatcher-ELP-GFP can be assembled into a multifunctional building block for GPV by the spontaneous formation of an isopeptide bond. mCherry-ZE-SpyTag and SpyCatcher-ELP-GFP were mixed at room temperature in 1xPBS buffer to produce mCherry-ZE-SpyTag/SpyCatcher-ELP-GFP (BB2) with an 80% yield after 30 minutes (Figure S5). Next, BB2 was mixed with ZR-ELP (60 μM) in a 1 M NaCl solution at two molar ratios (χ=0.01 and 0.05). After mixing the recombinant proteins on ice, the mixtures were incubated for 1 hour at 25□°C.

At χ=0.01, BB2 transitioned mostly into isolated GPVs (Figure 4 and S6), although some vesicle aggregation was observed. In contrast, even though phase separation of the building blocks occurred at χ=0.05, only coacervates and coacervate clusters were observed. During BB2 GPV assembly, a mCherry-ZE-SpyTag/SpyCatcher-ELP-GFP – ZR-ELP heterodimer can be formed. In that case, a portion of the hydrophobic ELP domain interacts with the coiled-coil domain of ZR-ELP homodimers and the SpyCatcher/SpyTag domain of BB2. Steric hindrance has been shown to destabilize bilayer formation in surfactant systems toward disordered or micelle-like aggregates, including coacervates.^51^

**Figure 4.**
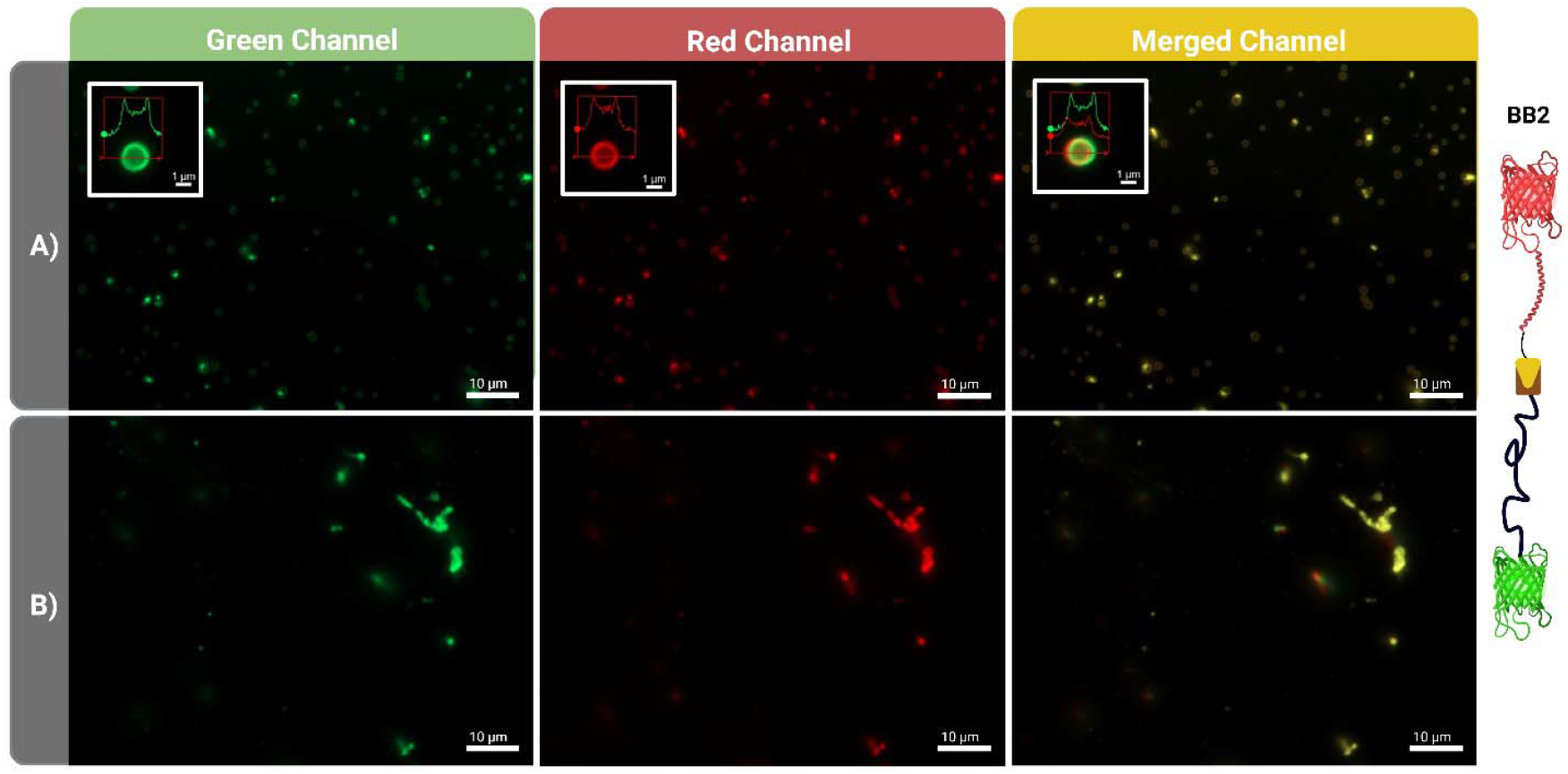
Epifluorescence micrographs of BB2-derived GPVs assembled with ZR-ELP. Vesicles were formed by incubating BB1 with ZR-ELP (60 μM) in 1 M NaCl for 1 hour at 25⍰°C. (A) χ = 0.01 molar ratio (BB1:ZR-ELP), (B) χ = 0.05.

### A single-unit ELP multifunctional bolaamphiphile: BB3

BB3 (mCherry-ELP-GFP) is the simplest of the three BBs and is composed of a near-symmetrical ELP bolaamphiphile. BB3’s lack of a coil domain enables us to examine the impact of incorporating globular proteins on GPV membranes without ZE/ZR interaction (Figure 1B). Since SpyCatcher-ELP-GFP bolaamphiphile (60 μM), undergoes phase separation at 1M NaCl and χ=1, forming coacervates, we suspected that it could transition into GPVs when mixed with ZR-ELP (Figure S2 and S7A). Indeed, when the χ ratio was reduced to 0.05 under the same salt conditions, the system transitioned into GPVs (Figure S7B). This observation aligns with prior work by Jang *et al*., who showed that the molar ratio of fusion proteins critically governs GPV self-assembly.^40^ We hypothesized that a recombinant triblock bolaamphiphile mCherry-ELP-GFP could be incorporated into the vesicle’s membrane when mixed with ZR-ELP above the critical salt concentration reported for eGFP-ZE GPVs, 0.91 M.

At χ=0.01, the mixture of BB3 and ZR-ELP fully transitioned into GPVs (Figure 5). We observed the same behavior as in BB1 and BB2 GPVs at this molar ratio (Figure S8). The GFP and mCherry domains have a high degree of similarity in size and structure. Thus, we expected the orientation of fluorescent proteins in the vesicle domain to be random. The formation of GPV clusters may be attributed to the surface display of GFP, which is known to weakly dimerize.^52^

**Figure 5.**
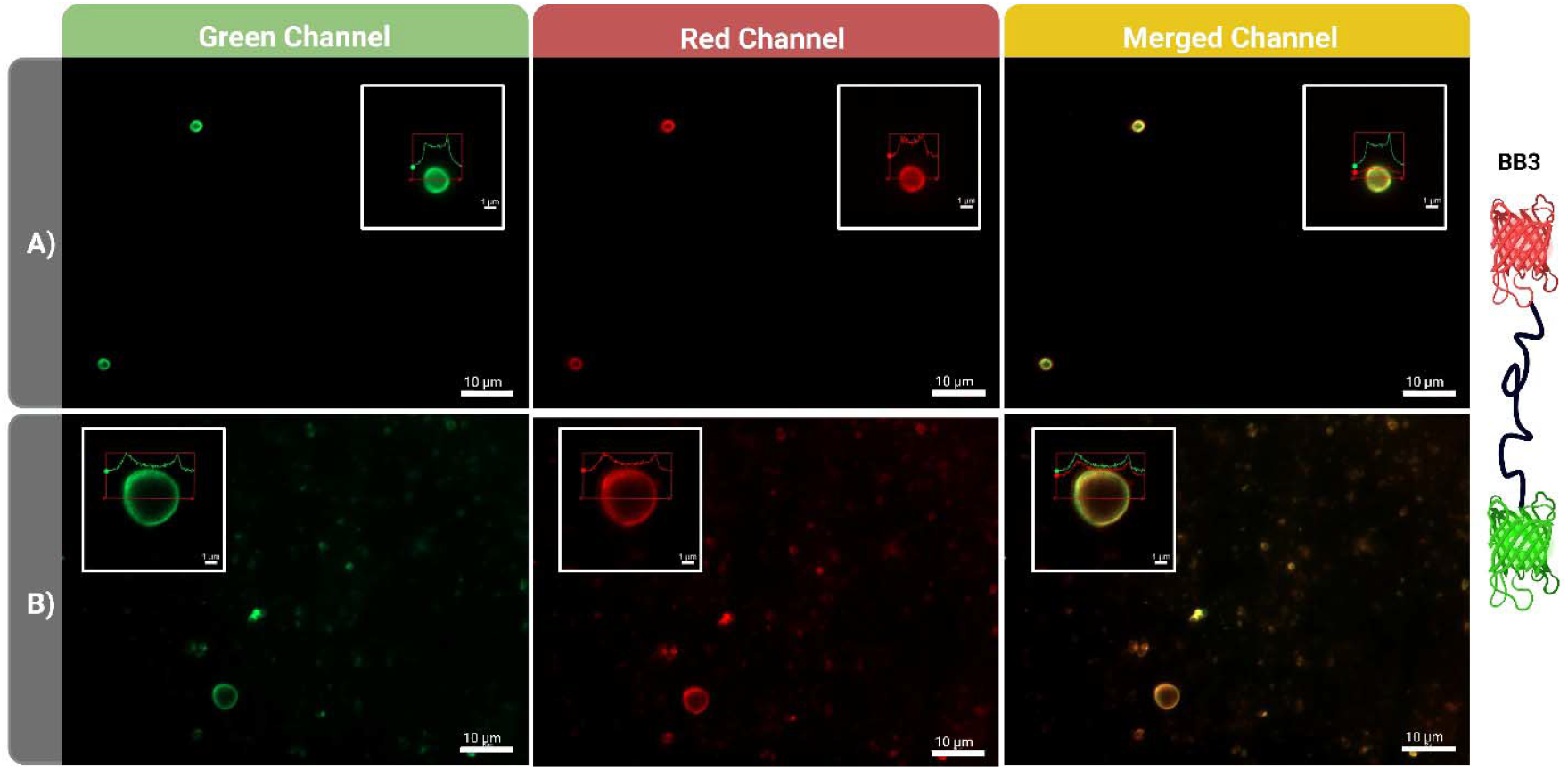
Epifluorescence micrographs of BB3-derived GPVs assembled with ZR-ELP. Vesicles were formed by incubating BB1 with ZR-ELP (60 μM) in 1 M NaCl for 1 hour at 25⍰°C. (A) χ = 0.01 molar ratio (BB1:ZR-ELP), (B) χ = 0.05.

At χ = 0.05, BB3 formed hollow vesicles of varying sizes. Unlike BB2, where increasing the BB:ZR-ELP molar ratio impaired vesicle assembly, BB3 was still able to form GPVs under the same conditions. One distinguishing feature of BB3 is its lack of a coil domain, whereas BB1 and BB2 include domains enriched in glutamic acid and/or arginine. Christensen *et al*. previously modeled the transition temperature of ELP fusion proteins as a function of the solvent-exposed amino acids in the globular domains.^46^ Their findings indicated that glutamic acid and arginine exert a strong influence on ELP transition temperature, with their solvent surface accessibility positively correlating with increased T_t_. This correlation may explain why BB3 retains the ability to form vesicles at χ = 0.05, whereas BB2 does not. Although BB3 lacks the modularity of the other designs, it offers a streamlined architecture for vesicle formation. These results suggest that even minimal bolaamphiphilic constructs can support GPV assembly, provided the BB:ZR-ELP molar ratio remains within an optimal range (χ = 0.01–0.05).

One limitation of this study was our inability to determine the orientation of bolaamphiphiles within GPVs, following methods successfully developed for liposomes.^53^ Based on the architecture of BB1 and BB2, strong parallel ZE/ZR interactions are expected to localize GFP exclusively within the vesicle lumen. However, attempts to experimentally verify this using antibody-based detection were unsuccessful due to key issues: the nonspecific association of antibodies with GPV membranes and the cross-reactivity of commercial anti-GFP and anti-mCherry antibodies. Furthermore, the semi-permeable nature of the GPV membrane (with a <40 kDa cutoff) allowed a highly specific anti-GFP nanobody to diffuse into the lumen, confounding efforts to distinguish between lumenal and membrane-associated GFP (Figures S10 and S11). Protease-based strategies also proved technically challenging due to issues such as the reassociation of cleaved protein domains on the GPV membrane. We are currently designing new approaches to reliably determine protein orientation in bolaamphiphile-based GPVs.

### Biophysical characterization of bolaamphiphilic GPVs

Bolaamphiphilic GPVs were characterized using dynamic light scattering (DLS) and atomic force microscopy (AFM) to quantify their size distribution and mechanical properties, parameters commonly reported in studies of GPVs.^2,6,13,40^ For BB1-derived GPVs, vesicle clustering at χ=0.01 ratio, prevented accurate determination of the hydrodynamic diameter. The samples presented high polydispersity, with aggregates forming either large or sedimenting particles, resulting in high cumulant fit and multimodal fit errors, as well as a low in-range figure (79%) (Figure S9). At χ=0.05, the vesicles exhibited a narrow size distribution, with an average hydrodynamic diameter (D_h_) of 727 nm (Figures S9A). In contrast, BB2- derived GPVs displayed a larger average D_h_ of 1.2 μm. For BB3 GPVs at χ=0.01, the DLS correlogram exhibited two decay regimes, and the resulting size distribution did not align with either the Z-average or microscopy observations (Figures S5). AFM imaging revealed vesicles with diameters approaching 4 μm (χ=0.01), likely explaining sedimentation at longer timescales (Figure 6). At χ=0.05, DLS measurements of BB3 GPVs were of poor quality, showing a high polydispersity index (PDI) and large multimodal fit error, indicative of significant heterogeneity and oversized aggregates (Figure S9).

**Figure 6.**
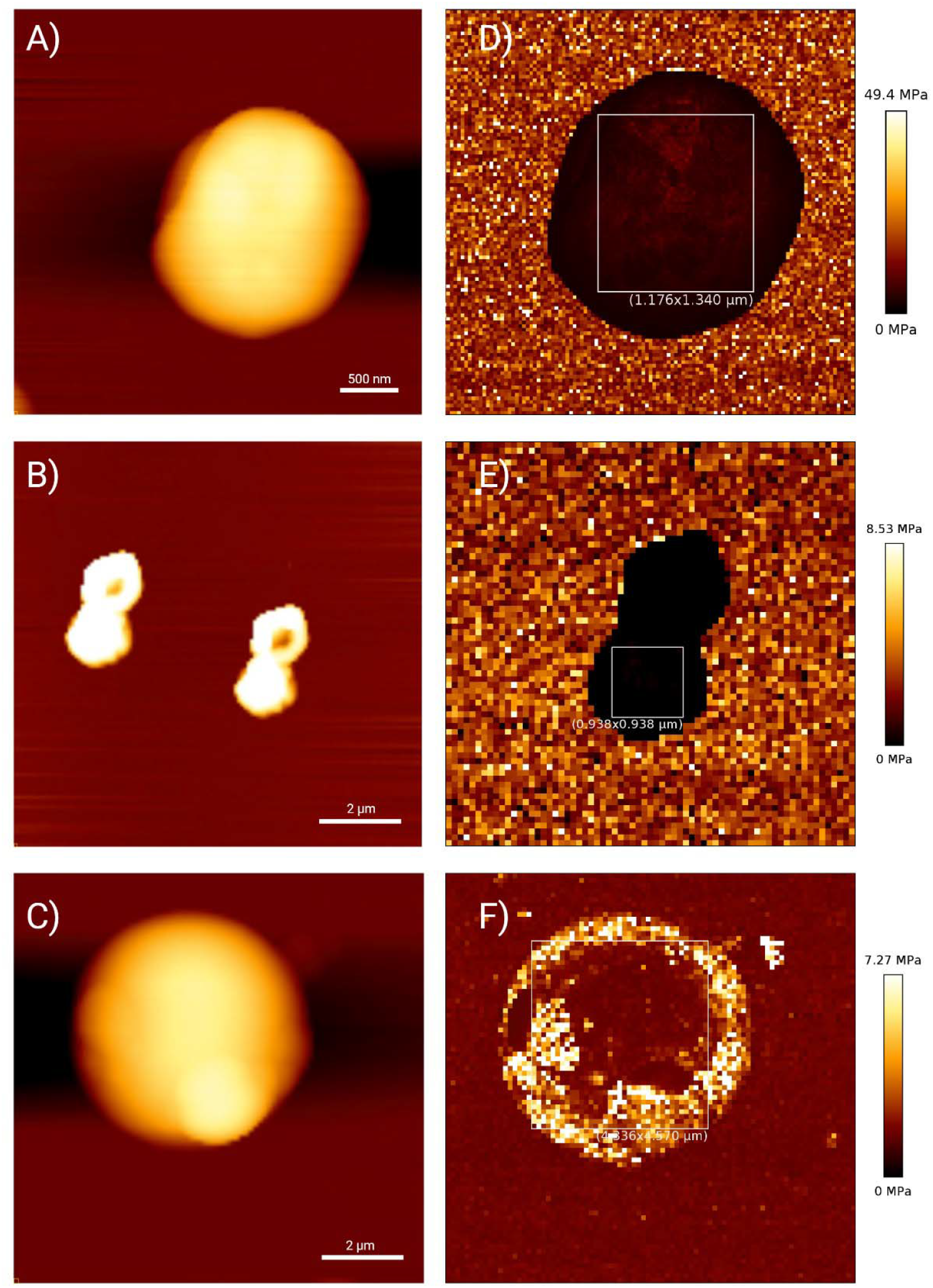
AFM characterization of GPVs assembled with BB1, BB2, and BB3. (A–C) Topographical AFM height plots for BB1, BB2, and BB3 GPVs, respectively, assembled at 60 μM ZR-ELP, 1 M NaCl, and χ = 0.01.(D–F) Young’s modulus maps of corresponding GPV membranes. The boxed regions in each map indicate the area used for statistical analysis.

Using force–deformation curves, we quantified the Young’s modulus of multifunctional GPV membranes assembled from BB1, BB2, and BB3 (Figure S12). The measured average moduli were 2.86 MPa (BB1), 1.26 MPa (BB2), and 1.63 MPa (BB3), slightly higher than the sub-MPa range of modulus of GPVs mainly made from mCherry-ZE (Figure 6).^1^ Notably, our vesicles were assembled under different conditions (1 M NaCl, 60 μM ZR-ELP, χ = 0.01), compared to the mCherry-ZE vesicles (0.3 M NaCl, 120 μM ZR-ELP, χ = 0.05), indicating that mechanical performance comparable to high protein content systems can be achieved using bolaamphiphilic designs at lower protein concentrations (Figure S13). This mechanical enhancement is likely driven by the combined effect of elevated ionic strength and the presence of ELP bolaamphiphiles.

Increased salt concentration is known to reduce the transition temperature (T□) of ELPs, facilitating denser molecular packing and reduced membrane hydration.^54–56^ These factors contribute to the formation of more cohesive and mechanically stiff vesicle membranes. The ability to preserve mechanical robustness while incorporating multiple functional domains in a modular format is particularly advantageous for material efficiency. Overall, the stiffness of these vesicles under high-salt conditions supports their suitability as environmentally tunable and shear-resistant nanocarriers for applications such as drug delivery and synthetic cellular systems.^57,58^

## Conclusions

This study presents a modular approach to building GPVs using ELP bolaamphiphiles. We developed a streamlined Golden Gate cloning strategy that facilitated the efficient assembly of three bolaamphiphilic building blocks (BB1–BB3), each varying in complexity and domain architecture. The constructs enabled precise genetic encoding and post-translational assembly of fusion proteins, incorporating two globular domains per bolaamphiphile—a design advancement over previously reported GPV systems, which relied on strong ZE/ZR dimer formation and were typically limited to the integration of a single globular domain.^2,4–8,28,39,40^ Our results highlight the versatility of this modular platform in supporting the systematic integration of functional proteins into synthetic vesicle membranes.

By varying the molar ratio (χ) of bolaamphiphiles to ZR-ELP, we observed differences in vesicle formation, morphology, and aggregation behavior, depending on the structural complexity of the building block. Fluorescence microscopy confirmed the successful incorporation of fluorescent domains, and GPV formation was consistently achieved under high-salt (1 M NaCl) conditions. The high modularity of BB1 offers flexibility in the architecture of the building blocks. Previous studies have shown that globular protein fusions containing ZE can be easily purified.^4^ Therefore, a wide range of functional bolaamphiphilic blocks with this structure could be assembled in GPVs. The site-specific conjugation of the modules guarantees accurate control over the building block composition. BB2 GPVs reduce the number of recombinant proteins to create a modular building block. Therefore, the assembly steps are simplified, and the cost of expressing a third recombinant protein is removed. Still, the flexibility to specifically bioconjugate easy-to-express recombinant proteins, incorporating multiple globular domains, is maintained. The simplicity of the BB3 construct, which lacks both ZE/ZR and SpyTag/SpyCatcher interactions, further demonstrates that a minimal triblock design is sufficient to mediate stable GPV formation, provided stoichiometric and ionic conditions are optimized. Previous bolaamphiphile systems, such as ZR-ELP-ZR constructs, were limited by their symmetric architectures, which restricted control over post-translational functionalization.^28^ By enabling modular, orthogonal assembly, our approach overcomes these constraints and broadens the design space for multifunctional synthetic vesicles.

Beyond demonstrating successful GPV formation, this study provides a generalizable approach for designing synthetic vesicles with customizable functionalities. By leveraging our cloning framework, additional functional proteins, enzymes, or transmembrane domains can be incorporated to expand GPV applications in biosensing, synthetic biology, and drug delivery. The modularity of this system enables further optimization of vesicle size, stability, and biochemical activity, paving the way for rationally designed bio-inspired materials. Importantly, we demonstrate that these bolaamphiphiles can self-assemble into GPVs with tunable mechanical properties, exhibiting Young’s moduli ranging from 1.26 to 2.86 MPa under 1 M salt conditions. This modulus range positions our GPVs as mechanically resilient yet

responsive compartments, suitable for physiological applications where membrane integrity under stress is critical. While this study does not yet achieve active transmembrane signal transduction, it establishes critical groundwork by demonstrating the programmable assembly of multifunctional GPVs. Future work will focus on integrating ligand-responsive receptor domains to achieve dynamic transmembrane signaling and coupling with intracellular actuators, advancing toward fully synthetic signal transduction in artificial cells.

## Methods

### Plasmid construction

The *pQE*9 expression vector with an N-terminal His□ tag was obtained from Qiagen (Catalog #: 32915). To eliminate a BsaI restriction site located within the ampicillin resistance gene, a silent point mutation was introduced by site-directed mutagenesis using primers RLEL007 (5’-gtgagcgtggctctcgcggta-3’) and RLEL008 (5’-cggctccagatttatcagc-3’), purchased from Sigma Millipore. This mutation preserved the encoded amino acid sequence while disrupting the BsaI recognition site. KOD One polymerase (Sigma Millipore Catalog #: KMM201NV), was used in all PCR reactions.

To generate the modified plasmid *pQE9-10xHis-2xBsaI*, four additional histidine codons and two BsaI sites were inserted via PCR. This was achieved using primers RLEL035 (5’- gcggcgttcagagaccgtcgacctgcagggtctccctgacttaattagctgagcttg-3’) and RLEL036 (5’- cgccctgaaaatacaggttttcggatccgtgatgatggtggtgatggtgatggtgatg-3’), purchased from Sigma Millipore. The PCR product replaced the original His-tag region, producing a plasmid with an extended 10xHis tag and two internal BsaI sites for modular cloning.

The receiver plasmid *pQE9-SpyCatcher-2xBsaI* was constructed using HiFi DNA assembly. A linear gene fragment encoding SpyCatcher003 with two BsaI sites was synthesized by Twist Bioscience. The *pQE9- 10xHis-2xBsaI* plasmid was digested with BamHI (NEB Catalog #: R3136S)and BsaI (NEB Catalog #: R3733S), and the 3.4 kb backbone fragment was purified via agarose gel electrophoresis and extracted using the QIAquick Gel Extraction Kit (Qiagen Catalog #: 28704). This purified backbone was assembled with the SpyCatcher003 insert using HiFi DNA Assembly Master Mix (NEB Catalog #: E2621S).

To generate *pQE9-mCherry-2xBsaI*, a similar HiFi DNA assembly approach was used. The linearized plasmid backbone was PCR-amplified from *pQE9-SpyCatcher-2xBsaI* using primers RLEL055 (5’- cctcctcgcccttgctcaccatctggaaatataagttttctgttgtg-3’) and RLEL056 (5’- gtcgaagcttcgcggtagtggatccgttcagagaccgtcgacctgcagggtc-3’), purchased from Sigma Millipore. The mCherry fragment was amplified from *pQE60-mCherry-ZE* using primers RLEL057 (5’- cacaacagaaaacttatatttccagatggtgagcaagggcgaggagg-3’) and RLEL058 (5’- gaccctgcaggtcgacggtctctgaacggatccactaccgcgaagcttcgac-3’), purchased from Sigma Millipore. The two fragments were assembled to yield the final mCherry construct with dual BsaI sites.

The gene encoding the ELP(VPGXG) □ □ polypeptide was divided into three segments and synthesized by Twist Bioscience. A double-stranded DNA fragment encoding superfolder GFP (sfGFP) was also purchased from Twist. These fragments were assembled into the *pQE9-SpyCatcher-2xBsaI* receiver plasmid using Golden Gate assembly, enabling the insertion of modular ELP and reporter elements. T7 DNA ligase (NEB Catalog #: M0318S), T4 DNA ligase reaction buffer (NEB Catalog #: B0202S), and BsaI (NEB Catalog #: R3733S) were used to set up the reaction.

The plasmid *pQE60-mCherry-ZE-SpyTag* was constructed by PCR using primers RLEL003 (5’- ggatgcgtataaaccgaccaaaggaggatcccatcaccatcaccatcactaagc-3’) and RLEL004 (5’- accatcacaatatgcgcagaaccgccaccaccagatctcagcggaccgtaac-3’), purchased from Sigma Millipore, with *pQE60-mCherry-ZE* as the template. The resulting PCR product was treated with the KLD enzyme mix (NEB Catalog #: M0554S) to circularize the plasmid and remove template DNA. The product was transformed into NEB Turbo chemically competent *E. coli* cells (NEB Catalog #: C2984H).

### Protein expression and purification

The plasmids containing the recombinant proteins in this work were transformed in BL21(DE)3 competent cells (NEB Catalog #: C2527H). Cells carrying *pQE*9 plasmid constructs were grown separately at 37°C in Lysogeny broth (LB) media with Ampicillin (GOLDBIO Catalog #: A-301-5) (100 mg/L) for 24 hours. Cultures to express proteins contained in *pQE*60 and *pET*29b (+) were grown separately at 37°C in Lysogeny broth (LB) media with ampicillin (100 mg/L) and Kanamycin (GOLDBIO Catalog #: K-120-5) (50 mg/L), respectively. When the optical density of the solution at 600 nm (OD_600_) reached 0.6-0.8, protein expression was induced with Isopropyl-β-thiogalactoside (IPTG, 1.0 mM) (GOLDBIO Catalog #: I2481C). Cells were pelleted by centrifugation after 4 hours and resuspended in lysis buffer. Liquid affinity chromatography was done in 10 mL Pierce^TM^ centrifuge columns (ThermoFisher Scientific Catalog #: 29924) using Nickel Agarose Beads (high density) (GOLDBIO Catalog #: H-350-100). Native conditions using phosphate-buffered saline (PBS) (Fisher Scientific Catalog #: BP39920) and imidazole (GOLDBIO Catalog #: I-902-25) buffers were used to purify all the proteins except ZR-ELP. Denaturing buffers previously reported were prepared for ZR-ELP purification. ^5^ ZR-ELP was dialyzed into 10 mM Tris-HCl (GOLDBIO Catalog #: T-400-10) and deionized water, and PBS 1x was the dialysis buffer for the fusion proteins containing globular domains. The purity of the proteins was assessed by sodium dodecyl sulfate-polyacrylamide gel electrophoresis (SDS-PAGE).

### Turbidity Assay

The protein solutions (1.5 ml) were prepared, transferred into cuvettes at 4 °C, and placed in the UV-2600 (Shimadzu). The turbidity of the protein solutions was assessed by measuring the optical density of transmitted light at 400 nm, a wavelength at which protein absorption is negligible. Changes in turbidity were monitored by recording the optical density every minute for 53 minutes in the temperature interval of 7-60°C.

### Fluorescence microscopy

The morphology of the protein vesicles was examined using epifluorescence and confocal microscopy with a 100× oil immersion objective lens (Axio Observer 7, LSM700, Carl Zeiss). The hydrodynamic diameter (D_h_) of the self-assembled vesicles was then determined using dynamic light scattering (DLS) with a Zetasizer Nano ZS (Malvern Instruments). The DLS instrument featured a 4 mW He–Ne laser operating at a wavelength of 633 nm and a detection angle of 173°, with measurements performed at 25°C. For DLS analysis, 40 μL samples were prepared in micro cuvettes.

### Atomic force microscopy

AFM samples were prepared using mica substrates (Ted Pella, INC.). A muscovite mica sheet (1 cm x 2 cm) was glued to a microscope slide and exfoliated. The exfoliated mica was pretreated with 100 µL of 125mg/mL Mg2+ in 1X PBS for 10 min, and the solution was removed. 50 µL of sample was then applied to the mica substrate and incubated for 5 min. The sample was then rinsed to remove non-adhered vesicles with 500 µL of 0.3M PBS thrice. 1 mL of 0.3M PBS was then applied to the sample after placing the glass slide on the AFM stage. All AFM measurements were taken using a JPK Nano wizard 4 XP (Bruker) and PFQNM-LC-V2 calibrated probes (Bruker). All images were captured at room temperature, in 1M PBS. Moduli were calculated in the JPK data processing software using extend force curves, applying the Hertz/Sneddon method, and assuming a spherical tip shape.

### Statistical analysis

All statistical analyses were conducted using a nonparametric approach due to the non-normal distribution of modulus values. A Kruskal–Wallis test assessed overall differences among vesicles (BB1, BB2, BB3, mCherry-ZE). To identify specific pairwise differences, Dunn’s post hoc test with multiple comparison correction was performed. Adjusted *p*-values were reported, and differences were considered significant at *p* < 0.05. Statistical significance is annotated in plots using letters: groups not sharing a letter are significantly different.

## Funding information

Grants from the National Science Foundation (2123592) supported this work.

## Notes

### Competing Interest Statement

The authors have declared no competing interest.

