## Supplemental File for "Rational Development of Recombinant ELP Bolaamphiphiles for the Controlled Construction of Multifunctionalized Globular Protein Vesicles"

**Supplemental Figures and Tables**

**Supplemental Figure 1.** SDS-PAGE analysis of the purified recombinant proteins used for BB1–BB3 vesicle construction

**Supplemental Figure 2.** Thermoresponsive phase transition of SpyCatcher-ELP-GFP and mCherry-ELP-GFP fusion proteins

**Supplemental Figure 3.** SDS-PAGE analysis of spontaneous ligation between ZR-SpyTag and SpyCatcher-ELP-GFP

**Supplemental Figure 4.** Fluorescent micrographs of GPVs formed by BB1 (0.01 or 0.05 molar ratio) and ZR-ELP (60 µM) in 1 M NaCl

**Supplemental Figure 5.** SDS-PAGE gel showing ligation of mCherry-ZE-SpyTag and SpyCatcher-ELP-GFP.

**Supplemental Figure 6.** Fluorescent micrographs showing phase transitions of GPVs assembled with BB2 and ZR-ELP at a 0.01 molar ratio

**Supplemental Figure 7.** Fluorescent micrographs of the phase transition of SpyCatcher-ELP-GFP and ZR-ELP

**Supplemental Figure 8.** Fluorescent micrographs showing phase transitions of GPVs assembled with BB3 and ZR-ELP at a 0.01 molar ratio

**Supplemental Figure 9.** Dynamic light scattering (DLS) analysis of GPVs assembled with BB1, BB2, and BB3 under varying molar ratios

**Supplemental Figure 10.** Assessment of protein orientation in GFP-ZE GPVs

**Supplemental Figure 11.** Evaluation of protein orientation in mCherry-ZE GPVs

**Supplemental Figure 12.** Mechanical characterization of GPVs assembled from BB1, BB2, and BB3

**Supplemental Figure 13.** Bar plot showing the mean ± SD of Young’s modulus for vesicles assembled from BB1, BB2, BB3, and mCherry-ZE fusion proteins

**Supplemental Table 1.** Amino acid sequence of the recombinant proteins

**Supplemental Table 2**. Plasmid utilized for cloning and/or expressing the building block elements

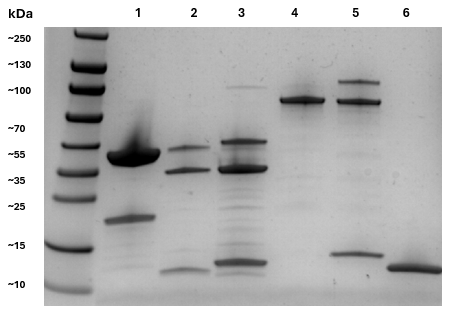
**Supplemental Fig. 1.** SDS-PAGE analysis of the purified recombinant proteins used for BB1–BB3 vesicle construction. Lane 1: ZR-ELP; Lane 2: mCherry-ZE; Lane 3: mCherry-ZE-SpyTag; Lane 4: SpyCatcher-ELP-GFP; Lane 5: mCherry-ELP-GFP; Lane 6: ZR-SpyTag.

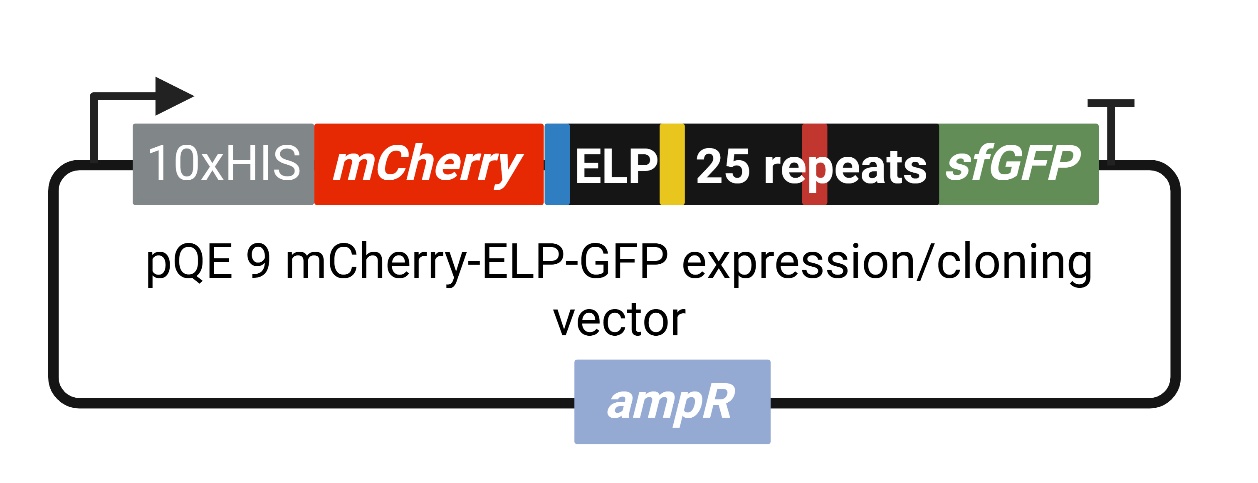

**Supplemental Fig. 2.** Expression plasmid for mCherry-ELP-sfGFP. The pQE9 mCherry-2xBsaI plasmid was used as the backbone. A Golden Gate assembly was set up with 4 inserts (the ELP gene split into three DNA fragments and the sfGFP gene fragment).

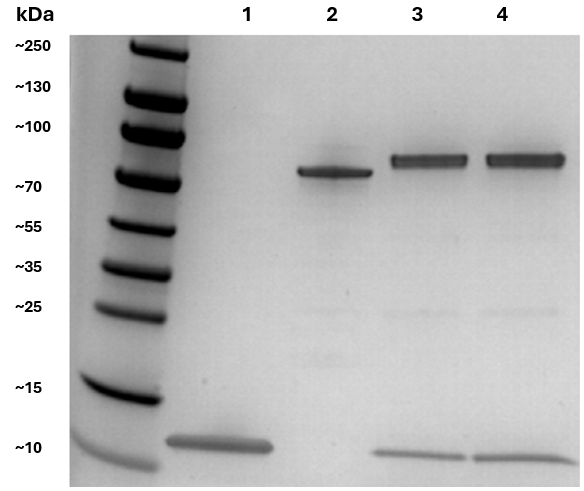
**Supplemental Fig. 3.** SDS-PAGE analysis of spontaneous ligation between ZR-SpyTag and SpyCatcher-ELP-GFP. Lanes: 1 – ZR-SpyTag; 2 – SpyCatcher-ELP-GFP; 3 – Ligated product at t = 0; 4 – Ligated product after one h.

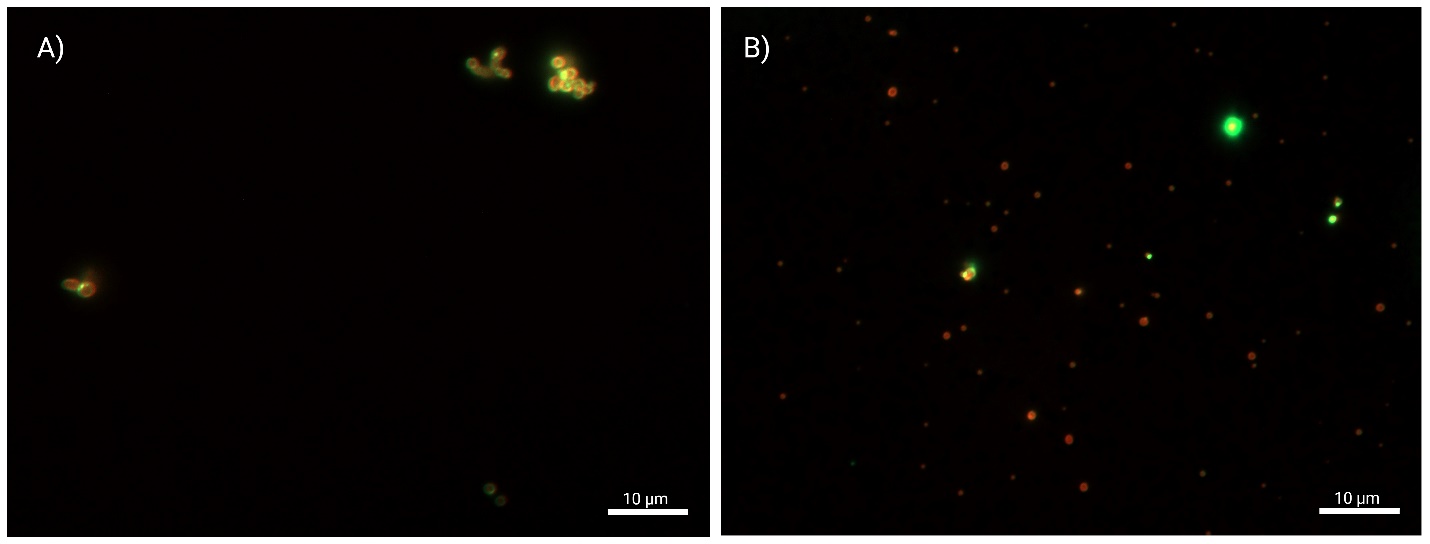
**Supplemental Figure 4.** Fluorescent micrographs of GPVs formed by BB1 and ZR-ELP (60 µM) in 1 M NaCl. Green (GFP) and red (mCherry) channels are merged. A) χ=0.01, B) χ=0.05. Scale bar = 10 µm.

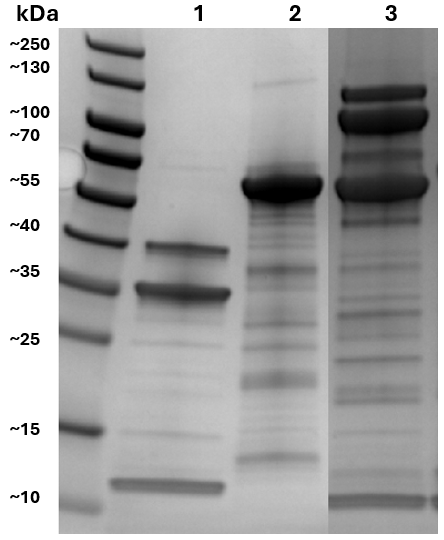
**Supplemental Figure 5.** SDS-PAGE gel showing ligation of mCherry-ZE-SpyTag and SpyCatcher-ELP-GFP. Lane 1: mCherry-ZE-SpyTag; Lane 2: SpyCatcher-ELP-GFP; Lane 3: Ligation product after one h. The ligation product shows two new bands at the expected molecular weight of the mCherry-ZE-SpyTag-SpyCatcher-ELP-GFP.

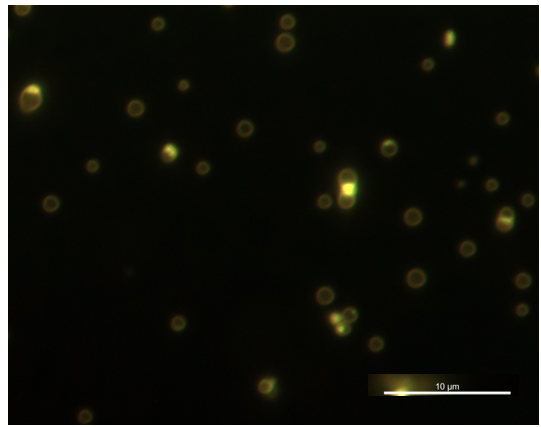
**Supplemental Figure 6.** Fluorescent micrographs showing phase transitions of GPVs assembled with BB2 and ZR-ELP at a 0.01 molar ratio (60 µM ZR-ELP, 1 M NaCl, 25 °C, 1 h). Scale bars = 10 µm.

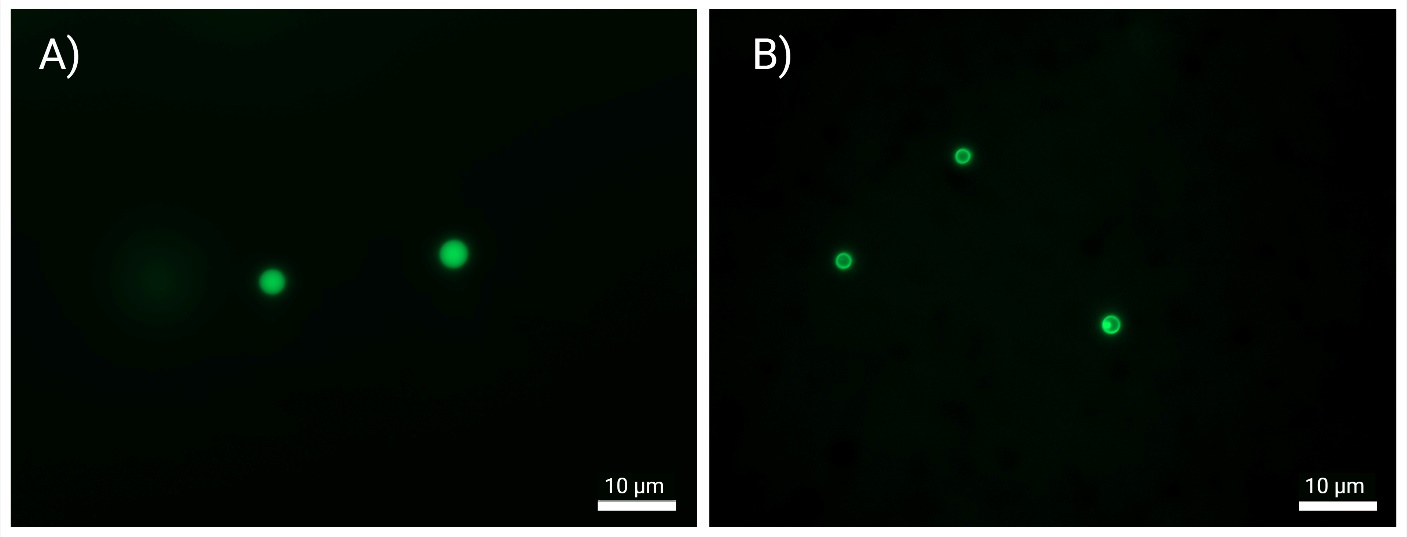
**Supplemental Figure 7.** Fluorescent micrographs of the phase transition of SpyCatcher-ELP-GFP and ZR-ELP. The concentration of NaCl was fixed at 1 M for protein assembly and phase transition at 25 ^0^C for 1 hour. A) SpyCatcher-ELP-GFP (120 µM), χ=1 B) SpyCatcher-ELP-GFP (6 µM), χ=0.05. The images show the green channel.

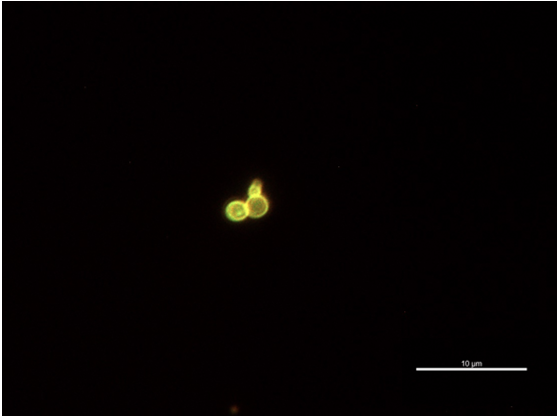
**Supplemental Figure 8.** Fluorescent micrographs showing phase transitions of GPVs assembled with BB3 and ZR-ELP at a 0.01 molar ratio (60 µM ZR-ELP, 1 M NaCl, 25 °C, 1 h).

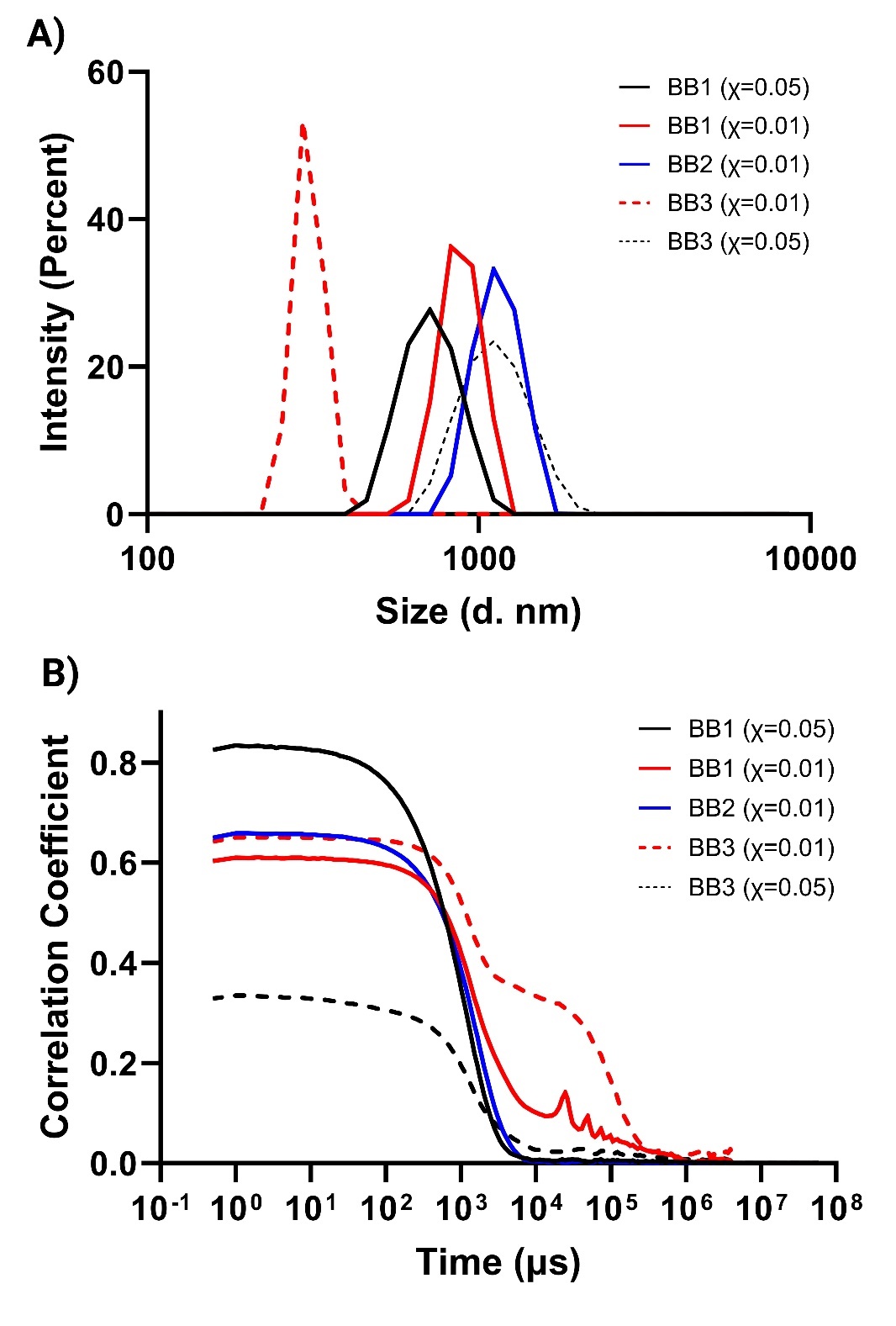
**Supplemental Figure 9.** Dynamic light scattering (DLS) analysis of GPVs assembled with BB1, BB2, and BB3 under varying molar ratios. All assemblies were prepared with 60 µM ZR-ELP and 1 M NaCl. (A) Intensity-weighted size distributions show that BB1- and BB2-containing GPVs (χ = 0.01 or 0.05) , and BB3 (χ = 0.05). (B) Correlogram plots of the corresponding samples show consistent differences in particle dynamics.

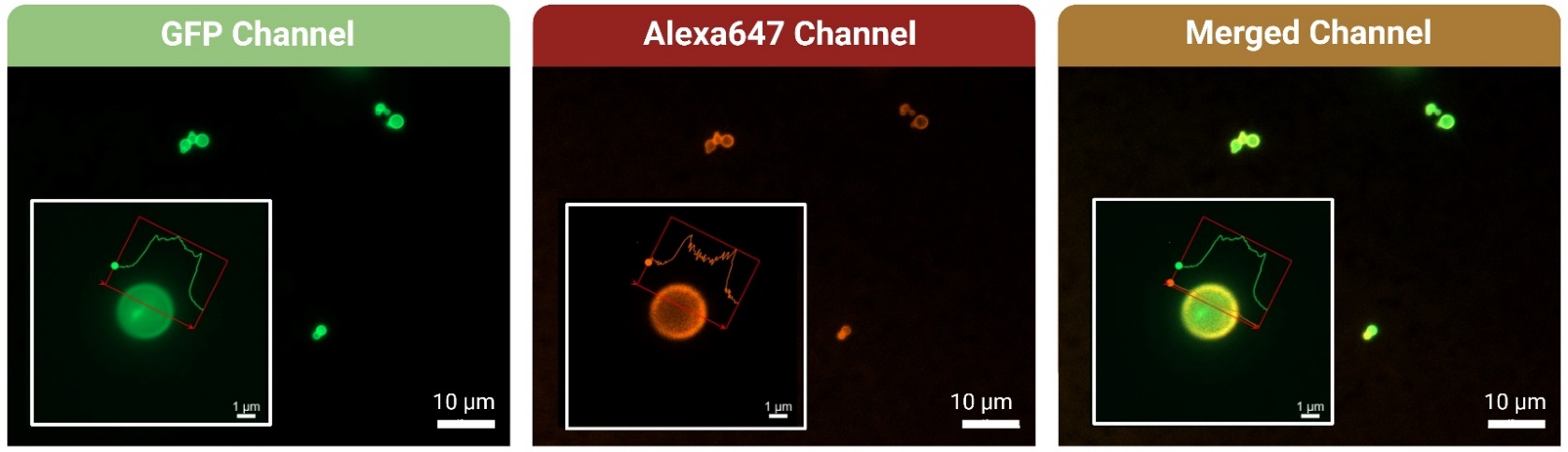
**Supplemental Figure 10.** Assessment of protein orientation in GFP-ZE GPVs. Vesicles were assembled on ice by mixing GFP-ZE (3 µM), ZR-ELP (60 µM), and NaCl (1 M), followed by incubation on ice for 15 minutes and subsequent incubation at 25 °C for 1 hour. To assess the orientation of GFP, an anti-GFP nanobody conjugated to Alexa Fluor® 647 (GFP-Booster Alexa Fluor® 647, 0.03 µM) was added to the vesicle solution, and samples were imaged using a fluorescence microscope (Axio Observer 7, LSM700, Carl Zeiss). Based on previous reports indicating external display of GFP in GFP-ZE vesicles, we hypothesized that the nanobody would label the vesicle membrane. Co-localization of Alexa Fluor® 647 and GFP fluorescence confirmed that GFP is displayed on the vesicle exterior.

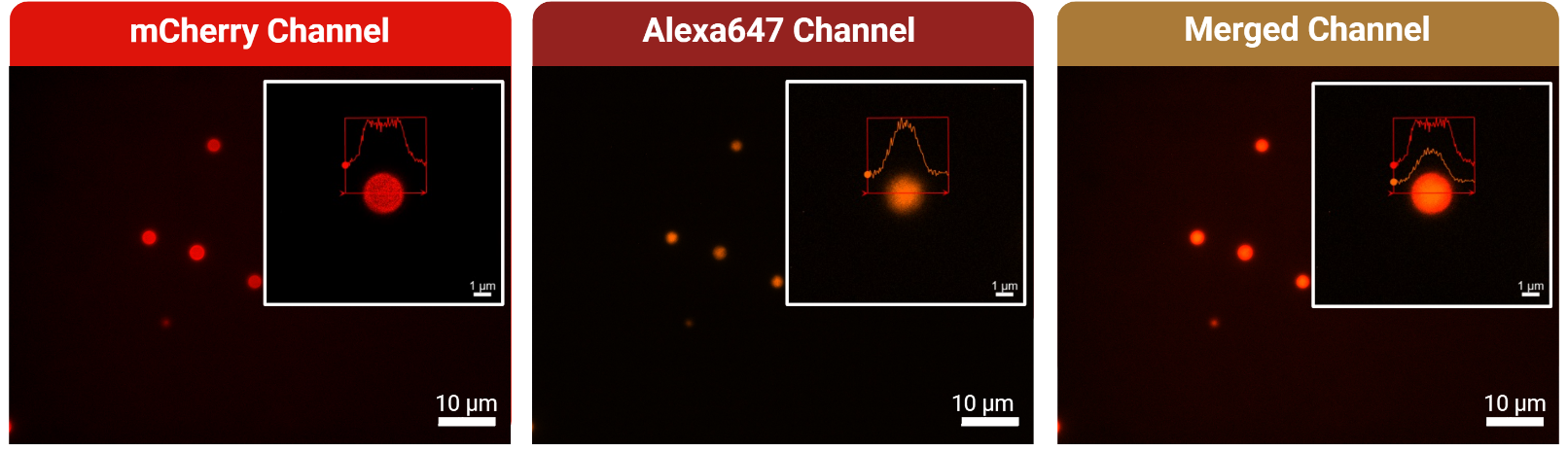
**Supplemental Figure 11.** Evaluation of protein orientation in mCherry-ZE GPVs. Vesicles were assembled by mixing mCherry-ZE (3 µM), ZR-ELP (60 µM), and NaCl (1 M) on ice for 15 minutes, followed by incubation at 25 °C for 1 hour. To evaluate protein orientation, an anti-GFP nanobody conjugated to Alexa Fluor® 647 (GFP-Booster Alexa Fluor® 647, 0.03 µM) was added to the vesicle solution, and samples were imaged using a fluorescence microscope (Axio Observer 7, LSM700, Carl Zeiss). Based on previous reports suggesting that mCherry-ZE single-layer GPVs are semipermeable (10 < cutoff < 40 kDa), we hypothesized that the nanobody would not permeate the vesicles or label the membrane. Contrary to our hypothesis, nanobody fluorescence was observed inside the vesicles, indicating that this approach cannot reliably determine protein orientation in GPV membranes.

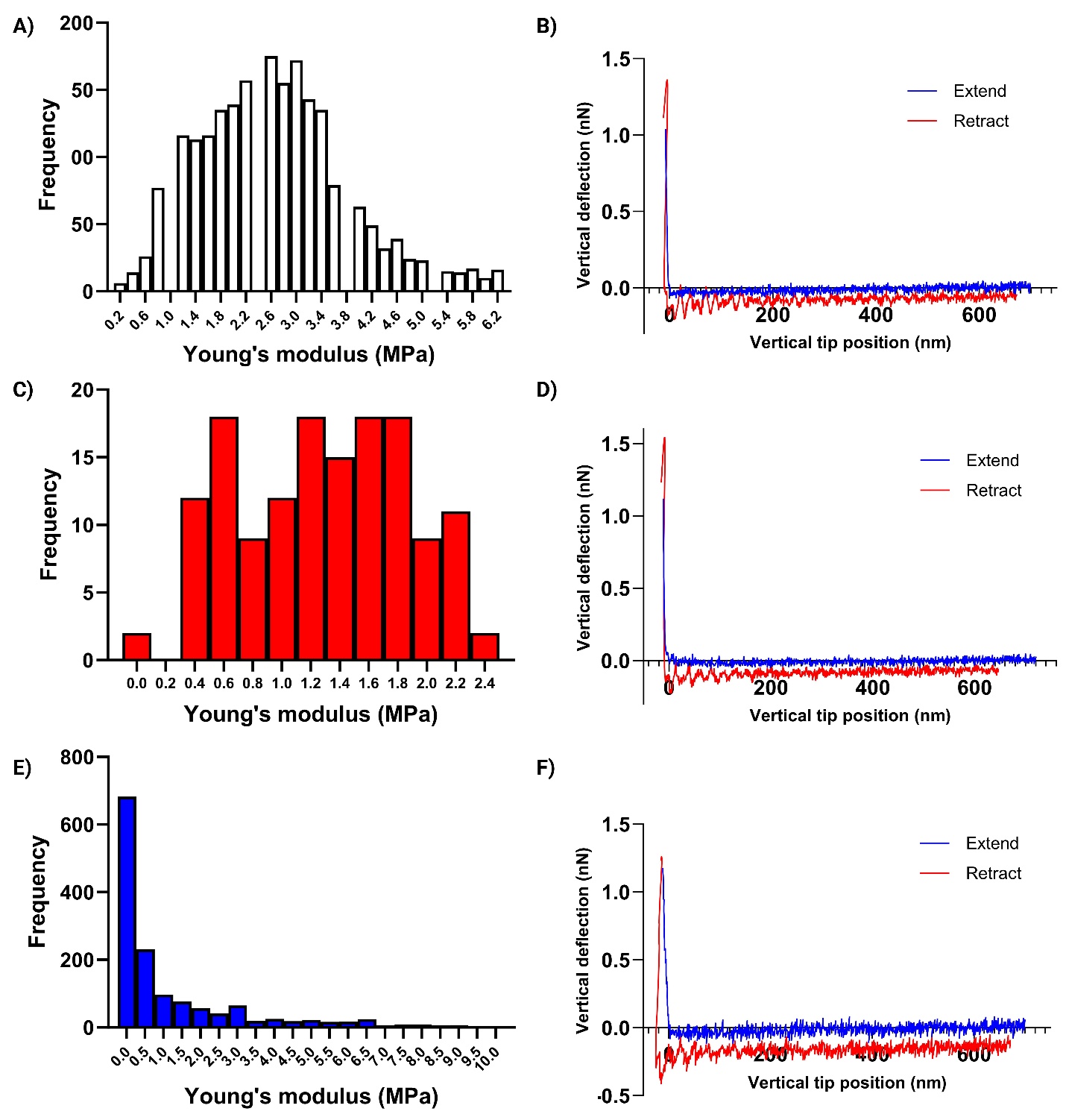
**Supplemental Figure 12.** Mechanical characterization of GPVs assembled from BB1, BB2, and BB3. A,C, E: Young’s modulus histograms from AFM measurements. B,D,F: Representative force-deformation curves.

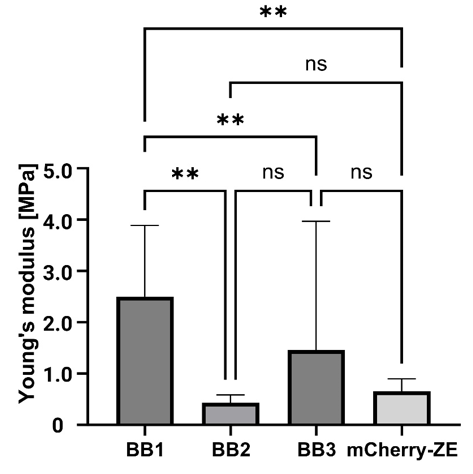
**Supplemental Figure 13.** Bar plot showing the mean ± SD of Young’s modulus for vesicles assembled from BB1, BB2, BB3, and mCherry-ZE fusion proteins. Mechanical properties were quantified from AFM force-deformation curves. Statistical analysis was performed using a Kruskal–Wallis test followed by Dunn’s multiple comparisons test (GraphPad Prism). BB1-derived vesicles exhibit significantly higher stiffness compared to BB2, BB3, and mCherry-ZE (**P** < 0.0001 and **P** = 0.0016, respectively), while no significant differences were observed among the other groups (**: significant, ns: not significant). BB1, BB2, and BB3 vesicles were assembled by incubating each building block with 60 μM ZR-ELP at a molar ratio of χ = 0.01 in 1 M NaCl at 25 °C for 1 hour. In contrast, mCherry-ZE vesicles were assembled under different conditions: 120 μM ZR-ELP, χ = 0.05, and 0.3 M NaCl.

**Supplemental Table 1**. The amino acid sequence of the recombinant proteins that comprise the molecular toolbox for assembling multifunctionalized GPVs.

| Recombinant protein | Amino acid sequence |
| --- | --- |
| mCherry-ZE | MGGSRSMVSKGEEDNMAIIKEFMRFKVHMEGSVNGHEFEIEGEGEGRPYEGTQTAKLKVTKGGPLPFAWDILSPQFMYGSKAYVKHPADIPDYLKLSFPEGFKWERVMNFEDGGVVTVTQDSSLQDGEFIYKVKLRGTNFPSDGPVMQKKTMGWEASSERMYPEDGALKGEIKQRLKLKDGGHYDAEVKTTYKAKKPVQLPGAYNVNIKLDITSHNEDYTIVEQYERAEGRHSTGGMDELYKSKLRGSGSLEIEAAALEQENTALETEVAELEQEVQRLENIVSQYRTRYGPLRSHHHHHH |
| ZR-ELP | MKGSLEIRAAALRRRNTALRTRVAELRQRVQRLRNEVSQYETRYGPLGGGGSGGGGSGVPGVGVPGVGVPGFGVPGVGVPGVGVPGVGVPGVGVPGFGVPGVGVPGVGVPGVGVPGVGVPGFGVPGVGVPGVGVPGVGVPGVGVPGFGVPGVGVPGVGVPGVGVPGVGVPGFGVPGVGVPGVGVPGC |
| mCherry-ZE-SpyTag | MGGSRSMVSKGEEDNMAIIKEFMRFKVHMEGSVNGHEFEIEGEGEGRPYEGTQTAKLKVTKGGPLPFAWDILSPQFMYGSKAYVKHPADIPDYLKLSFPEGFKWERVMNFEDGGVVTVTQDSSLQDGEFIYKVKLRGTNFPSDGPVMQKKTMGWEASSERMYPEDGALKGEIKQRLKLKDGGHYDAEVKTTYKAKKPVQLPGAYNVNIKLDITSHNEDYTIVEQYERAEGRHSTGGMDELYKSKLRGSGSLEIEAAALEQENTALETEVAELEQEVQRLENIVSQYRTRYGPLRSGGGGSAHIVMVDAYKPTKGGSHHHHHH |
| SpyCatcher003-ELP-sfGFP | MRGSHHHHHHHHHHGSDYDIPTTENLYFQGAMVTTLSSLSGEQGPSGDMTTEEDSATHIKFSKRDEDGRELAGATMELRDSSGKTISTWISDGHVKDFYLYPGKYTFVETAAPDGYEVATPIEFTVNEDGQVTVDGEATEGDAHTGSSGSVPGVGVPGVGVPGFGVPGVGVPGVGVPGVGVPGVGVPGFGVPGVGVPGVGVPGVGVPGVGVPGFGVPGVGVPGVGVPGVGVPGVGVPGFGVPGVGVPGVGVPGVGVPGVGVPGFGVPGVGVPGVGVPGGGGSGSRKGEELFTGVVPILVELDGDVNGHKFSVRGEGEGDATNGKLTLKFICTTGKLPVPWPTLVTTLTYGVQCFARYPDHMKQHDFFKSAMPEGYVQERTISFKDDGTYKTRAEVKFEGDTLVNRIELKGIDFKEDGNILGHKLEYNFNSHNVYITADKQKNGIKANFKIRHNVEDGSVQLADHYQQNTPIGDGPVLLPDNHYLSTQSVLSKDPNEKRDHMVLLEFVTAAGITHGMDELYK |
| mCherry-ELP-GFP | MRGSHHHHHHHHHHGSDYDIPTTENLYFQMVSKGEEDNMAIIKEFMRFKVHMEGSVNGHEFEIEGEGEGRPYEGTQTAKLKVTKGGPLPFAWDILSPQFMYGSKAYVKHPADIPDYLKLSFPEGFKWERVMNFEDGGVVTVTQDSSLQDGEFIYKVKLRGTNFPSDGPVMQKKTMGWEASSERMYPEDGALKGEIKQRLKLKDGGHYDAEVKTTYKAKKPVQLPGAYNVNIKLDITSHNEDYTIVEQYERAEGRHSTGGMDELYKSKLRGSGSVPGVGVPGVGVPGFGVPGVGVPGVGVPGVGVPGVGVPGFGVPGVGVPGVGVPGVGVPGVGVPGFGVPGVGVPGVGVPGVGVPGVGVPGFGVPGVGVPGVGVPGVGVPGVGVPGFGVPGVGVPGVGVPGGGGSGSRKGEELFTGVVPILVELDGDVNGHKFSVRGEGEGDATNGKLTLKFICTTGKLPVPWPTLVTTLTYGVQCFARYPDHMKQHDFFKSAMPEGYVQERTISFKDDGTYKTRAEVKFEGDTLVNRIELKGIDFKEDGNILGHKLEYNFNSHNVYITADKQKNGIKANFKIRHNVEDGSVQLADHYQQNTPIGDGPVLLPDNHYLSTQSVLSKDPNEKRDHMVLLEFVTAAGITHGMDELYK* |
| ZR-SpyTag | MKGSLEIRAAALRRRNTALRTRVAELRQRVQRLRNEVSQYETRYGPLGGGGSGGGGSGAHIVMVDAYKPTK |

**Supplemental Table 2**. Plasmid utilized for cloning and/or expressing the building block elements.

| Plasmid name | Benchling file | Source |
| --- | --- | --- |
| *pQE9-10xHis-2xBsaI* | https://benchling.com/s/seq-Lc8g9V2fa2yDZNF2NwBs?m=slm-Cr3qq2OmvWdJwz10yNvr | This work |
| *pQE9-SpyCatcher-2xBsaI* | https://benchling.com/s/seq-gI7r5deY2NgwWr34OjXR?m=slm-L2g7xtTL09eWD3yXKcln | This work |
| *pQE9-mCherry-2xBsaI* | https://benchling.com/s/seq-Gml76uPgHnGfrs1KW6qM?m=slm-Cc9id4zAdn1jTjTVb1We | This work |
| *pQE60-mCherry-ZE* | https://benchling.com/s/seq-lkXl3SqT9Au1K27Fxo0N?m=slm-0hNDqWc26tYA03a7LOen |  |
| *pQE9-SpyCatcher-ELP-GFP* | https://benchling.com/s/seq-a4CNkOqalHZ8zZsb0ExQ?m=slm-3dzi3R6ssRVCGJq7HdYu | This work |
| *pQE9-mCherry-ELP-GFP* | https://benchling.com/s/seq-OnGO0bzpCbTrLVDaqx2r?m=slm-AupCG9nWIWHXi2n8FPqC | This work |
| *pQE60-mCherry-ZE-SpyTag* | https://benchling.com/s/seq-vLdsggOrGRZva5ge0LkA?m=slm-LRPHWx7Dh7ccMRqSZ8oG | This work |
| *pQE60-ZR-ELP* | https://benchling.com/s/seq-SUlEvwULI6kvl7UYS2vY?m=slm-joyKPiq91kNTSanMHxK3 |  |
